# Intracellular glutamine fluctuates with nitrogen availability and regulates *Mycobacterium smegmatis* biofilm formation

**DOI:** 10.1101/2025.06.18.660496

**Authors:** Elizabeth Varner, Mitchell Meyer, Jocelyn Whalen, Yu-Hao Wang, Carlos Rodriguez, Ifra Malik, Steven J. Mullett, Stacy L. Gelhaus, William H. DePas

## Abstract

Biofilm formation allows pathogenic nontuberculous mycobacteria (NTM) to adhere to household plumbing systems and has been observed *in vivo* during human infection. Glucose drives NTM aggregation *in vitro*, and ammonium inhibits it, but the regulatory systems controlling this early step in biofilm formation are not understood. Here, we show that a variety of carbon and nitrogen sources have similar impacts on aggregation in the model NTM *Mycobacterium smegmatis* as glucose and ammonium, suggesting that the response to these nutrients is general and likely sensed through downstream, integrated signals. Next, we performed a transposon screen in *M. smegmatis* to uncover these putative regulatory nodes. Our screen revealed that mutating specific genes in the purine and pyrimidine biosynthesis pathways caused an aggregation defect, but supplementing with adenosine and guanosine had no impact on aggregation either in a *purF* mutant or WT. Realizing that the only genes we hit in purine or pyrimidine biosynthesis were those that utilized glutamine as a nitrogen donor, we pivoted to the hypothesis that intracellular glutamine could be a nitrogen-responsive node affecting aggregation. We tested this hypothesis in defined M63 medium using targeted mass spectrometry. Indeed, intracellular glutamine increased with nitrogen availability and correlated with planktonic growth. Furthermore, a *garA* mutant, which has an artificially expanded glutamine pool in growth phase, grew solely as planktonic cells even without nitrogen supplementation. Altogether these results establish that intracellular glutamine controls *M. smegmatis* aggregation, and they introduce flux-dependent sensors as key components of the NTM biofilm regulatory system.

**Importance:** A subset of nontuberculous mycobacteria (NTM), including *Mycobacterium abscessus*, are opportunistic pathogens that can cause severe pulmonary infections. *M. abscessus* is present in showerhead biofilms, one of the reservoirs from which it can infect susceptible individuals. Moreover, *M. abscessus* can exist as biofilms during pulmonary infections, and biofilm formation *in vitro* renders *M. abscessus* more tolerant to antibiotics. The ability to inhibit NTM biofilm formation could therefore help us better prevent and treat NTM infections. However, the regulatory systems controlling NTM biofilm formation, which could include targets for anti-biofilm therapeutics, are poorly understood. The significance of this work is that it reveals intracellular glutamine as an important node controlling initiation of biofilm formation in the model NTM *Mycobacterium smegmatis*. Building on this foundation, future studies will investigate how NTM biofilms can be dispersed by altering glutamine levels and will describe how NTM translates intracellular glutamine to alteration of surface adhesins.

## Introduction

Biofilms confer bacterial cells with specific benefits such as surface adhesion and protection against external stressors(1), but they are also energetically costly to produce and can impose growth limitations on subpopulations within the biofilm(2, 3). Bacteria therefore have regulatory systems that only trigger biofilm assembly in certain environments, presumably where the benefits of living in multicellular structures outweigh the costs. Since bacteria occupy diverse niches, biofilm-permissive conditions, and the chemical cues that trigger biofilm formation, differ from one bacterial species to the next. For example, *Staphylococcus aureus* biofilm formation is typically enhanced by the presence of glucose(4), whereas *Escherichia coli* biofilm formation can be suppressed by glucose(5). *Streptoccocus mutans* biofilm formation is inhibited by oxygen(6), whereas *Campylobacter jejuni* forms more robust biofilms in aerobic conditions(7).

Similarly, biofilm regulatory systems differ substantially from species to species. Some species like *Pseudomonas aeruginosa* and *Vibrio cholerae* make extensive use of intracellular cyclic-di-GMP (c-di-GMP), which can respond to multiple extracellular stimuli, to control the transition between biofilm formation and motility(8, 9). Enterics like *Escherichia coli* and *Salmonella enterica* also utilize c-di-GMP, but it is just one of many inputs that control the master biofilm regulator CsgD(10, 11). The promoter region controlling expression of *csgD* is amongst the largest and most regulated intergenic regions in the genome, with multiple global and specific regulators binding overlapping regions(11). In *Bacillus subtilis*, various phosphotransferases integrate on a specific master biofilm regulator, Spo0A, a process that is also influenced by quorum sensing and ties in with sporulation(12). Despite their differences, these systems share the ability to collate multiple pieces of information from the extracellular environment and integrate them into a central regulatory pathway that controls whether bacteria initiate biofilm formation or not. Enough pro-biofilm signals ultimately lead to the production of matrix components such as polysaccharides, proteinaceous fibers, and eDNA.

Nontuberculous mycobacteria (NTM) are emerging and problematic pathogens that form biofilms during pulmonary infections and in the built environment(13–16). NTM biofilm formation is typical in that it renders cells more resistant to stresses such as antibiotic exposure and disinfectants(17, 18), but mycobacterial biofilms are unusual in at least two important ways. First, the mycobacterial biofilm matrix differs from that of canonical Gram-positive and Gram-negative bacteria. There are no well-described proteinaceous fibers in the NTM biofilm matrix, and the role of cellulose (present in some mycobacterial biofilms but with no clear synthetase) and eDNA (strain to strain variability) is unclear(19–23). Instead, the mycobacteriales-specific mycomembrane creates a hydrophobic cell surface environment in which lipophilic molecules such as glycopeptidolipids (GPLs) and long-chain mycolic acids can affect cell-to-cell adhesion during biofilm development(18). Secondly, while environmental cues such as iron and oxygen influence biofilm maturation(24, 25), there is no integrated regulatory system described for NTM that controls biofilm initiation. Contributing to our lack of insight into this potential regulatory system is the tendency of mycobacteria to clump in most liquid media(26–28). A true planktonic state is therefore not easily accessible *in vitro* as a biofilm comparator. Detergents are frequently added to liquid medium to physically force mycobacterial cells apart, but how growth in detergent affects the activity of potential biofilm regulatory systems is not well understood(18).

In this study, we sought insight into the regulatory processes controlling the first step in NTM biofilm formation; the transition from planktonic cells to aggregates. We build off recent results showing that NTM can grow planktonically without detergents when provided with relatively poor carbon and replete nitrogen sources(26). Guided by an unbiased transposon screen in the model NTM *Mycobacterium smegmatis*, we focused on metabolic nodes that respond to carbon and nitrogen availability that could be involved in regulating biofilm formation. We hit purine and pyrimidine biosynthesis genes multiple times in our screen, so we first explored aggregation dynamics of an *M. smegmatis* Δ*purF* mutant. Those experiments led us to the hypothesis that the utilization of the glutamine pool for purine/pyrimidine biosynthesis, and not purines and pyrimidines themselves, affects aggregation. Follow up experiments with WT and mutant strains in defined medium, coupled with metabolic analysis, supported this hypothesis. Altogether, this study shows that glutamine fluctuates with nitrogen availability in *M. smegmatis*, and it reveals intracellular glutamine as a key regulator of the planktonic/aggregation transition in NTM. More broadly, we introduce flux-dependent sensors, intermediates in central metabolism whose intracellular concentrations correlate with flux through specific metabolic pathways(29), as integrated nutrient-availability sensors from which NTM may control aggregation. These results build a foundation for future studies aimed at understanding the regulatory network controlling the initial decision of whether or not NTM form a biofilm in a given chemical environment.

## Results

### Multiple carbon and nitrogen sources impact *M. smegmatis* aggregation

*M. smegmatis* grows as aggregates and then spontaneously disperses in rich TYEM (tryptone, <u>y</u>east <u>e</u>xtract, <u>M</u>gSO_4_) medium (Fig. 1A)(26). Addition of glucose to TYEM rich medium extends and increases the *M. smegmatis* aggregation phase, and addition of ammonium leads to earlier dispersal and planktonic growth(26). Mycobacteria co-catabolize available carbon and nitrogen sources, with no obvious preference for any specific nutrient(30–32). We therefore began this study by testing whether aggregation and dispersal induction were specific to glucose and ammonium in rich medium. We supplemented TYEM with four different carbon sources, glucose, glycerol, pyruvate, and succinate, and found that all increased aggregation and delayed growth of planktonic cells, with some variation in the magnitude of the effect (Fig. 1B). Similarly, while ammonium was more efficient, nitrate supplementation also triggered dispersal (Fig. 1C). These results suggested to us that *M. smegmatis* likely does not key in on specific external nutrient sources to translate carbon/nitrogen availability into changes in aggregation. Instead, we hypothesized that *M. smegmatis* was sensing integrated metabolic nodes as a proxy for total carbon/nitrogen flux and was using those nodes to regulate aggregation.

**Figure 1.**
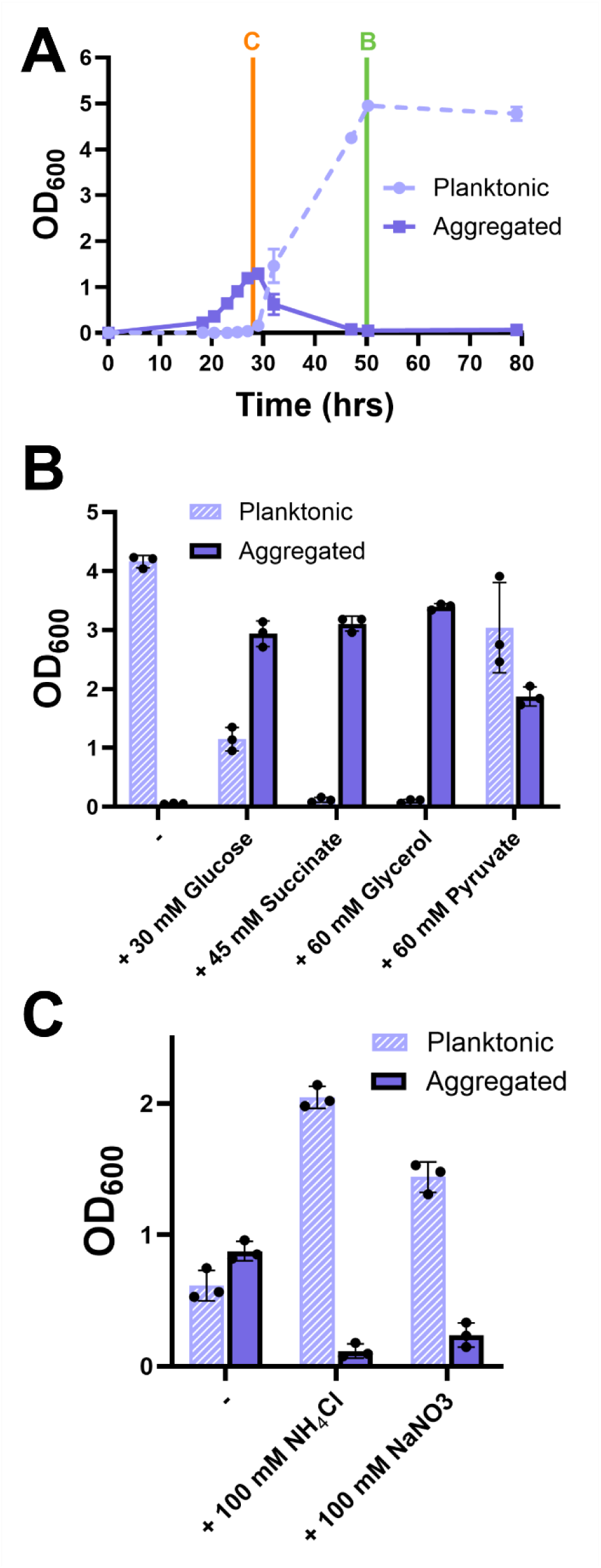
A) Aggregation dynamics of *M. smegmatis* in nutrient-rich medium (TYEM) – aggregates and planktonic cells were separated at indicated timepoints and the OD_600_ of each fraction was recorded. B) OD_600_ of planktonic and aggregate fractions of *M. smegmatis* grown in TYEM with indicated carbon supplements at 50 hours (timepoint indicated in A for reference). C) OD_600_ of planktonic and aggregate fractions of *M. smegmatis* grown with ammonium or nitrate supplements at 28 hours (timepoint indicated in A for reference). Error bars indicate mean ±SD (n=3). Dots indicate A) mean (n=3) or B) C) single replicates.

### Transposon screen to identify metabolic nodes controlling aggregation

To determine the identity of these putative regulatory nodes, we performed a transposon mutagenesis screen with phage phiMycoMarT7(33) in *M. smegmatis*. WT *M. smegmatis* wrinkles on TYEM agar plates (Fig. 2A), and wrinkling generally correlates with aggregation in liquid cultures(26). We therefore visually screened transductants for altered colony morphologies on TYEM agar plates in a primary screen, phenotyping at days 5 and 12 (Fig. 2B). We passaged candidates in liquid medium and then plated cell suspensions on new TYEM agar plates for a secondary screen (Fig. 2C). As expected, transductants with smooth colony morphotypes typically correlated with low aggregation in liquid medium, while hyper-wrinkling on agar plates correlated with constitutive aggregation (Fig. S1)(17, 26). Arbitrarily-primed PCR was performed on strains that differed from WT at days 5 and/or 12 in the secondary screen to determine where the transposon inserted in the genome(34, 35). In total, we generated a library of 133 independent transductants with an identified Tn landing site that showed altered colony morphology in our secondary screen (Table S1).

**Figure 2.**
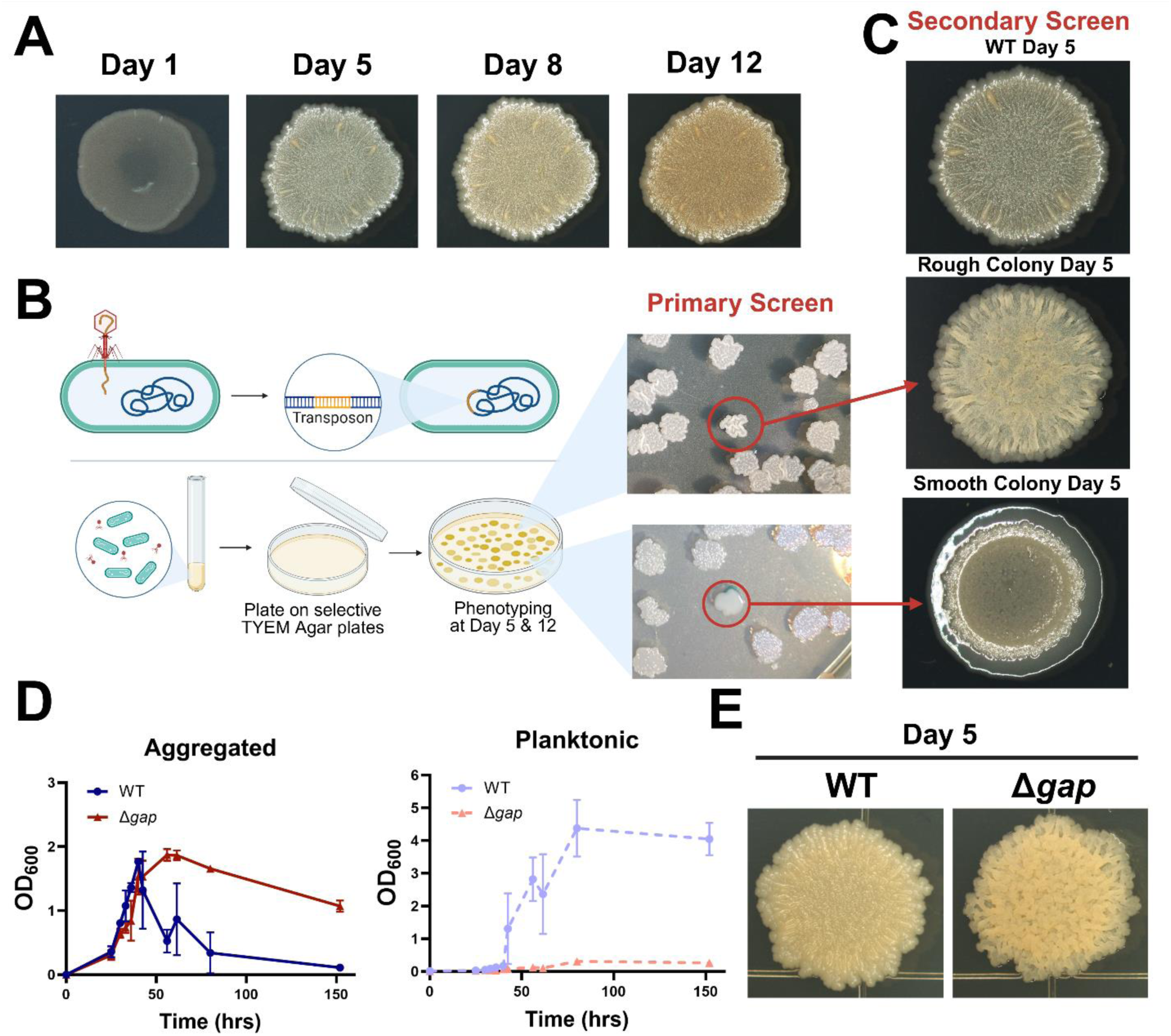
A) Colony morphology of WT *M. smegmatis* on TYEM agar plates. B) Cartoon depicting transposon mutagenesis screening and examples of candidate colonies on primary screen on TYEM agar plates. C) Colony morphology of example rough and smooth transductants compared to WT on Day 5 in secondary screen on TYEM agar plates. D) Aggregation dynamics of WT *M. smegmatis* and Δ*gap* in TYEM liquid medium. Error bars indicate mean ±SD (n=3). E) Colony morphology of WT vs. Δ*gap* on TYEM agar plates after 5 days of growth.

While the main purpose of our screen was to identify regulatory nodes that control aggregation in response to carbon or nitrogen availability, we also accrued several hits in genes known to code for surface molecules. For example, 15 of our 133 hits (>11%) were in genes that are involved in GPL biosynthesis (Table S1). *M. abscessus* frequently accumulates mutations in GPL biosynthesis genes during infection(36). These mutations result in dramatic ‘rough’ colony variants that constitutively aggregate in liquid medium(26). To verify that this association holds true in *M. smegmatis*, we knocked out *gap* (MSMEG_0403), a transport protein which is required for GPL transport to the surface(37). As expected, Δ*gap* constitutively aggregated in TYEM liquid medium and showed a rough colony morphology (Fig. 2D). In future studies, we will attempt to identify the surface adhesins/matrix components that change in response to carbon and/or nitrogen availability, and the putative and documented adhesins we hit in this screen will serve as candidates. For the purposes of this study, they serve as proof of principle, demonstrating that our screen can effectively identify genes that affect aggregation.

### Mutations in specific purine biosynthesis genes suppress *M. smegmatis* aggregation

Next, we manually reviewed our list, with a specific focus on genes involved in central metabolic pathways or carbon and/or nitrogen regulatory pathways. The first trend that stood out to us was a bevy of hits in purine and pyrimidine biosynthesis. We identified five mutants with a Tn landing site in purine biosynthesis genes (three hits in *purF*, one in *purL*, and one in *purQ*) and three mutants with a Tn landing site in pyrimidine biosynthesis genes (two hits in *carB* and one in *carA*) (Table S1). Both purine and pyrimidine biosynthesis are linked to biofilm regulation in a variety of bacterial species(38, 39), and carbon and nitrogen availability can influence purine and pyrimidine biosynthesis(40, 41). We therefore considered that intracellular purine or pyrimidine concentrations could fluctuate with carbon or nitrogen availability and have a direct role in *M. smegmatis* aggregation. Since we obtained more hits in purine biosynthesis, we focused on that pathway. To explicitly test how a defect in *purF* affects aggregation, we made a clean deletion of *purF* in *M. smegmatis* using recombineering(42). Δ*purF* showed a smooth colony morphotype, while the three *purF* transposon mutants showed intermediate phenotypes (Fig. 3A). Δ*purF* demonstrated an aggregation defect in TYEM liquid medium without demonstrating an obvious growth defect, likely because enough purines are available in the yeast extract for growth (Fig. 3B). Similarly, an *S. aureus* Δ*purF* mutant showed a growth defect in defined medium but not rich medium(43). Complementing with WT *purF* under control of the constitutive hsp60 promoter restored WT aggregation (Fig. S2). If a purine deficiency was the reason for a decrease in aggregation, we predicted that adding supplemental adenosine and guanosine to the media could restore aggregation in Δ*purF*. However, addition of these two purines to levels that can restore WT growth in an *M. smegmatis* Δ*purF* mutant in defined medium(44), did not affect aggregation (Fig. S3). Similarly, addition of adenosine and guanosine to WT did not affect colony morphology on TYEM agar plates (Fig. S4). Therefore, while our results suggested that purine biosynthesis can affect aggregation in *M. smegmatis*, they also suggested that the impact on aggregation was not directly due to accumulation of the end products, adenosine and guanosine. To find a viable path forward, we scrutinized the specific genes in the purine and pyrimidine biosynthesis pathways that we hit in our screen.

**Figure 3.**
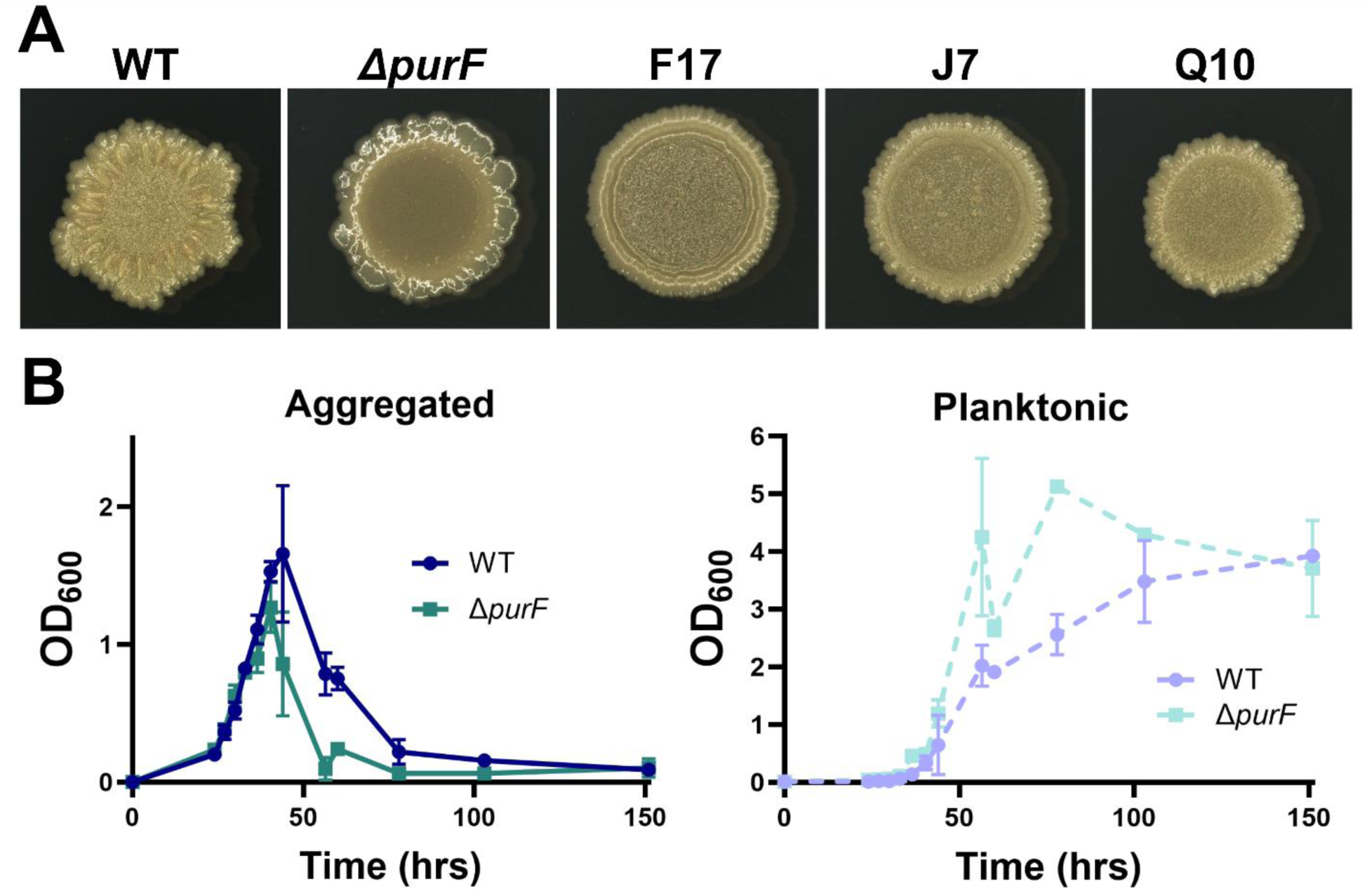
A) Colony morphology of WT *M. smegmatis*, *ΔpurF*, and the three transductants with Tn insertion sites in the *ΔpurF* gene at day 5 on TYEM agar plates. B) Aggregation dynamics of WT *M. smegmatis* and *ΔpurF* in TYEM. Error bars indicate mean ±SD (n=3).

### Screen hits cluster around enzymes utilizing glutamine for amination reactions

We soon realized that PurF and the PurLQS complex catalyze the only two amination reactions in purine biosynthesis, both of which utilize glutamine as a nitrogen donor. Strikingly, we did not hit any non-aminating enzymes in purine biosynthesis, of which there are many(45). Similarly, the only genes we hit in pyrimidine biosynthesis, *carA* and *carB*, produce a protein complex that catalyzes the only reaction in pyrimidine biosynthesis that utilizes glutamine as an amine donor (bicarbonate + gln + 2ATP → carbomyl-phosphate + glu + 2ADP + Pi). We did not hit any other genes in pyrimidine biosynthesis. Focusing in on enzymes that utilize glutamine as a nitrogen source, we realized that another of our hits was in a Glu-tRNA^Gln^ amidotransferase, which uses glutamine as a nitrogen donor to convert Glu-tRNA^Gln^ to Gln-tRNA^Gln^ (46). In all, nine out of 133 sequenced hits (>6%) mapped to genes coding for enzymes that use glutamine as the nitrogen source in an amination reaction (Fig. 4). Glutamine is central to bacterial nitrogen regulation; bacteria assimilate nitrogen from inorganic sources such as ammonium or nitrate chiefly through glutamine synthetase (glu + NH_3_ + ATP → gln + ADP + Pi) and glutamate dehydrogenase 1 (alpha-ketoglutarate (AKG) + NH_3_ + NADPH → glu + NADP^+^)(47–49). Glutamine and glutamate are then the amine donors for biosynthesis of all nitrogen-containing macromolecules in the cell(47–49). Glutamate is the nitrogen donor for most transaminase reactions (∼85% of nitrogenous compound biosynthesis, most amino acids) while glutamine donates nitrogen for purine and pyrimidine biosynthesis, glucosamine, and others (∼15% of nitrogenous compound biosynthesis)(48). It is glutamine, though, that commonly serves as an indicator of cellular nitrogen status for bacteria; the intracellular glutamine pool swells with increased nitrogen availability and shrinks in nitrogen deplete conditions(47, 49, 50). Most, if not all, bacterial regulators that respond to nitrogen availability sense, directly or indirectly, intracellular glutamine(47, 49).

**Figure 4.**
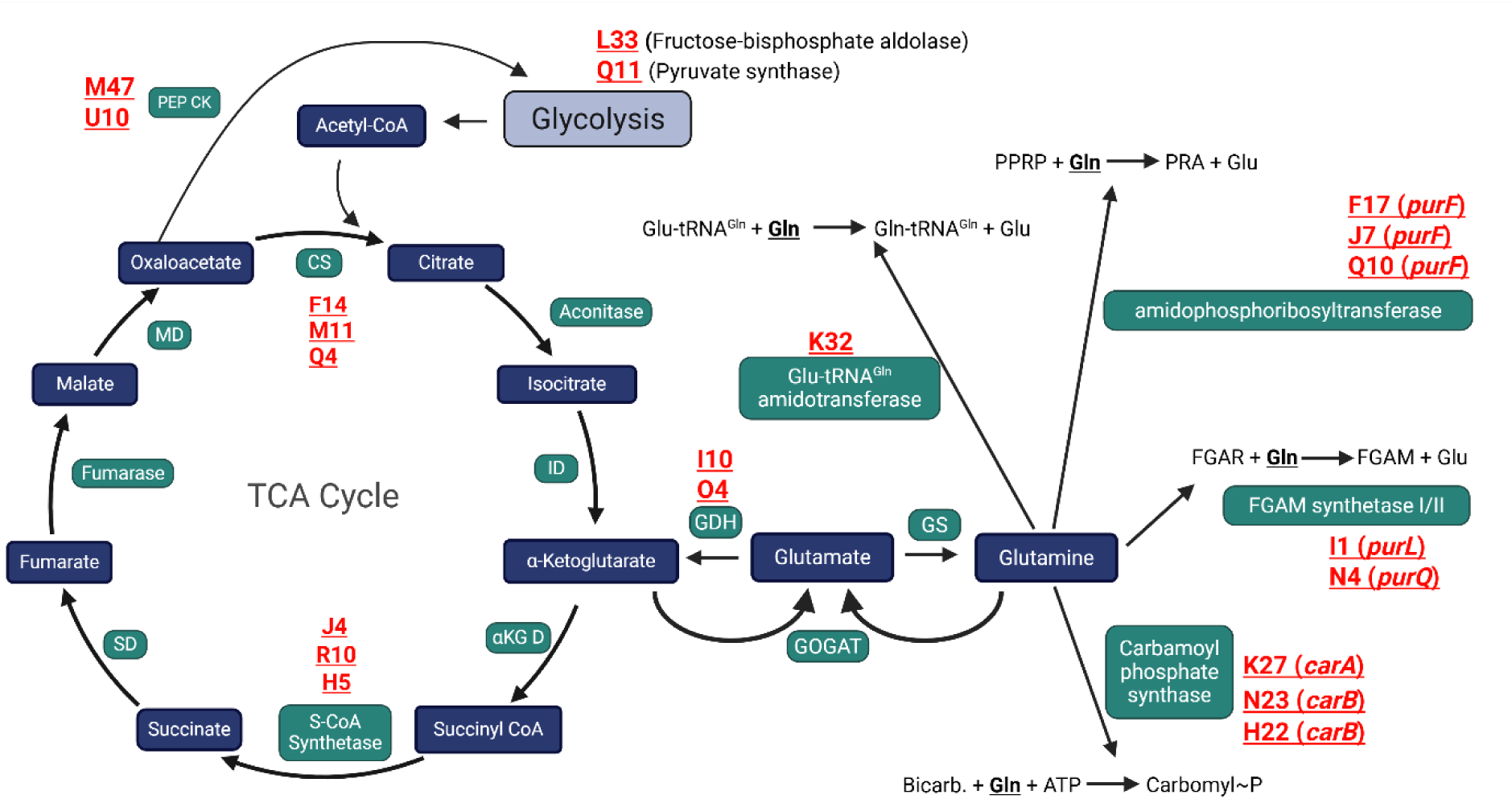
Schematic of the TCA cycle and enzymes hit in the transposon mutagenesis screen that utilize glutamine as a nitrogen donor. Blue boxes represent metabolites, green boxes are enzymes, and red text indicate independent transposon hits in the associated gene.

Glutamine levels rise in mycobacteria with ammonium availability(50), which suggests that higher intracellular glutamine would correlate with planktonic growth. If deletion of *purF* leads to a larger intracellular glutamine pool, since that pool is no longer being utilized for *de novo* purine biosynthesis, the larger glutamine pool in Δ*purF* would again correlate with more planktonic growth/earlier dispersal (Fig. 3A). Supplementing with adenosine or guanosine would not be expected to directly affect the glutamine pool, which could explain why they did not affect aggregation in Δ*purF* or WT (Fig. S3, S4). We therefore pivoted to the hypothesis that intracellular glutamine responds to nitrogen availability and regulates aggregation in *M. smegmatis*.

### Design of a defined medium to assess the role of glutamine in aggregation

To directly test for a correlation between nitrogen availability, intracellular glutamine, and decreased aggregation in *M. smegmatis*, we first designed a defined medium which would allow us to track aggregation, accurately tune carbon and nitrogen sources, and limit background signal for metabolomic analysis. M63 with pyruvate as the major carbon source and ammonium as the major nitrogen source supports *M. smegmatis* growth as planktonic cells. Conversely, M63 with glycerol as the carbon source and glutamate as the sole nitrogen source supports aggregative growth(26). Here we found that M63 with 20 mM glucose and 20 mM glutamate as the main carbon/nitrogen sources mirrors aggregation kinetics in rich medium, with distinct aggregation and dispersal phases (Fig. 5A). The addition of 20 mM NH_4_Cl to this recipe prevents aggregation and results in growth solely as planktonic cells (Fig. 5A). Colony biofilms developed on M63 agar plates, and colony morphology mirrored aggregation phenotypes, with WT *M. smegmatis* wrinkling more on plates without ammonium (Fig. 5B). To further verify that the cell-to-cell adhesion underlying liquid aggregation is important for biofilm development, we also assessed pellicle biofilms in M63 glucose +/-NH_4_Cl. Like the colony morphology phenotype, pellicles wrinkled more without ammonium (Fig. 5C). Altogether these data suggest that M63 glucose is a suitable defined medium for assessing the role of nitrogen in *M. smegmatis* biofilm formation.

**Figure 5.**
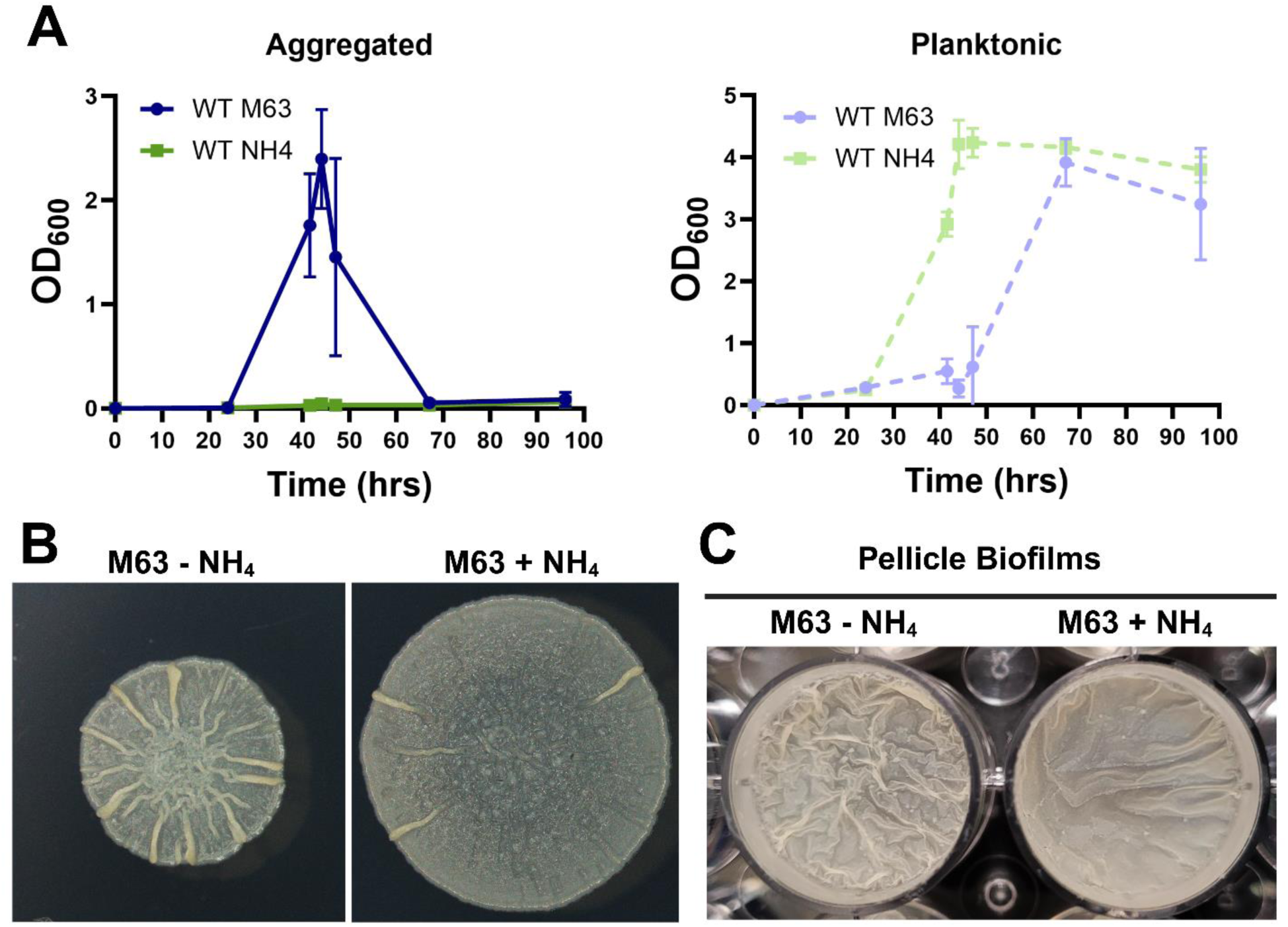
A) Aggregation dynamics of WT *M. smegmatis* in M63 glucose medium with and without ammonium. B) Colony morphology of WT *M. smegmatis* on M63 glucose agar plates with and without ammonium. C) Pellicle biofilms after 4 days of growth in M63 glucose medium with and without ammonium. Error bars indicate mean ±SD (n=3).

### Intracellular glutamine fluctuates with nitrogen availability and causes changes to aggregation

To test whether intracellular glutamine correlates with nitrogen availability in *M. smegmatis*, we harvested cells in log phase growth in either condition (glucose M63 +/-NH_4_Cl), extracted metabolites, and performed targeted liquid chromatography high resolution-mass spectrometry (LC-HRMS) to measure intracellular glutamine. As expected, the intracellular glutamine pool increased with NH_4_Cl supplementation, providing support for the hypothesis that there is a correlation between a larger glutamine pool and decreased aggregation (Fig. 6A, B, C). As a control we also tested for fluctuation in AKG levels, which can control cellular process in response to nitrogen availability in some species, but those levels did not change (Fig. S5)(49, 51, 52). Next, to probe a causal relationship between high intracellular glutamine and decreased aggregation, we sought to alter intracellular glutamine through genetic manipulation.

**Figure 6.**
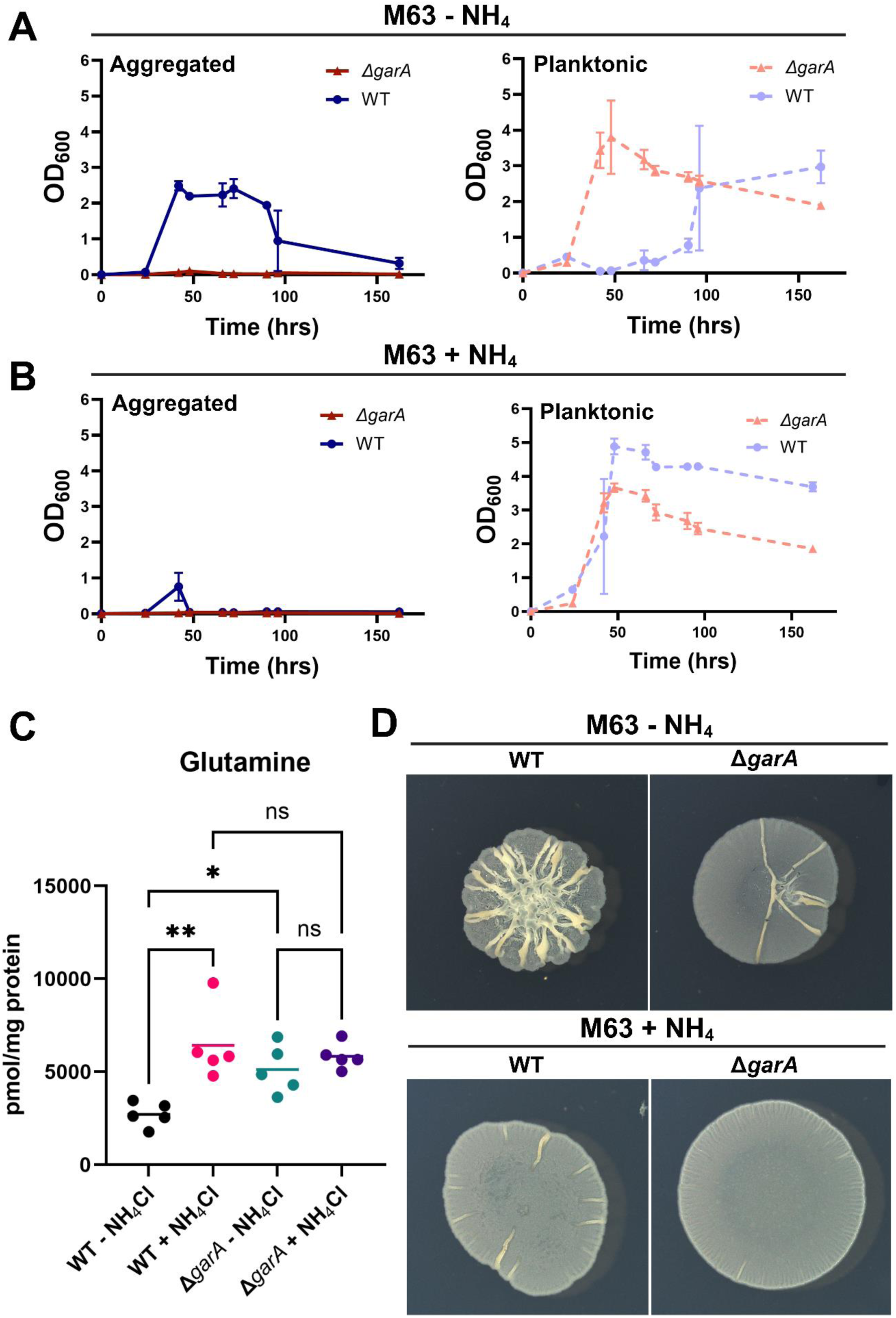
A) Aggregation dynamics of WT *M. smegmatis* and *ΔgarA* in M63 medium without ammonium. B) Aggregation dynamics of WT *M. smegmatis* and *ΔgarA* in liquid M63 medium with 20 mM ammonium. C) Intracellular glutamine quantification of WT *M. smegmatis* and *ΔgarA* in liquid M63 medium with and without ammonium at 40 hours. Lines indicate sample mean (n=5). D) Colony morphology of WT *M. smegmatis* and *ΔgarA* on M63 agar plates with and without ammonium. Error bars indicate mean ±SD (n=3).

We focused on GarA, a forkhead association domain-containing protein that binds and affects multiple enzymes in the AKG-glutamate-glutamine node based on its phosphorylation status(53, 54). Δ*garA* can grow in defined medium, and the net result of an *M. smegmatis* Δ*garA* mutant is a larger glutamine pool in growth phase(53). We therefore predicted that a Δ*garA* mutant would aggregate less than WT. Indeed, our Δ*garA* mutant demonstrated a high intracellular glutamine pool in M63 glucose, and grew solely as planktonic cells even without NH_4_Cl addition (Fig. 6A, B, C). Cloning *garA* behind its native promoter into the integration vector pMH94 rescued aggregation at early timepoints to a degree (Fig. S6). However, the aggregates were either too small or fragile to stay on the 10 µm filter, so we were unable to quantify this complementation phenotype. Colony morphology again mirrored aggregation phenotypes (Fig. 6D). Taken together our results support a model in which the intracellular glutamine pool fluctuates in response to nitrogen availability and restrains *M. smegmatis* aggregation.

## Discussion

NTM infections such as those caused by *M. abscessus* and *Mycobacterium avium* are very difficult to effectively treat with antibiotics(55–59). The long treatment durations required for NTM infections also lead to numerous undesirable side effects, including substantial ototoxicity from aminoglycosides(59, 60). Compounding the ineffectiveness of antibiotic treatments against NTM is the disconnect between *in vitro* drug susceptibility tests and clinical efficacy of any given drug(57, 59, 61, 62). We and others hypothesize that mechanisms of physiological tolerance, including biofilm formation, likely contribute to this disconnect(62). Biofilm dispersal agents, when applied in conjunction with antibiotics, can significantly improve treatment of bacterial infections in animal models(63). *M. abscessus* and *M. smegmatis* both aggregate and disperse *in vitro* and respond to carbon and nitrogen cues in rich medium similarly(26). By studying *M. smegmatis* biofilm regulation, we therefore hope to reveal pathways that can be targeted such that we can control NTM biofilm formation and dispersal, and subsequent antibiotic tolerance, *in vivo*.

Biofilm regulation in mycobacteria is poorly understood. A major barrier to describing putative biofilm regulatory systems is that mycobacteria constitutively aggregate in most liquid media, leading to the pervasive dogma that mycobacteria do not grow as free-living, planktonic cells(26–28). In a simple model of biofilm development, planktonic cells first adhere to one another and/or to a surface. Then, these aggregates further develop into a mature biofilm(64). Under the paradigm of constitutive mycobacterial aggregation, signals, genes, and gene products that are present in already formed NTM biofilms, or that are necessary for biofilm maturation, can be and have been probed and described. For example, a dedicated chaperone involved in mycolic acid biosynthesis, GroEL1, is necessary for *M. smegmatis* biofilm maturation(65). Glycopeptidolipids (GPLs) affect biofilm robustness and can be regulated by specific environmental cues such as oxygen(27, 66–68). Biofilms form in low iron and magnesium, but supplementation of these two nutrients allows for more robust biofilms in *M. smegmatis* and *M. abscessus*, respectively(24, 69). However, without an experimentally accessible planktonic state, the first step in biofilm formation, the transition between planktonic cells and aggregates, has not been well-studied in NTM and is poorly understood. The recent finding that carbon and nitrogen availability controls the planktonic:aggregate transition in *M. smegmatis* and *M. abscessus*(26) allows us to probe that node, with the expectation that it will reveal novel NTM regulatory systems and potential targets for anti-biofilm therapeutics.

It may be that a typical biofilm regulatory system exists in mycobacteria, but we have not been able to fully describe it due to the aforementioned difficulties in studying the planktonic:aggregate transition. However, the unusual aspects of mycobacterial aggregation, namely that mycomembrane components serve, at least in part, as the biofilm matrix, and that mycobacteria readily aggregate in most liquid media, force us to consider that biofilm regulation may be a fundamentally different process in mycobacteria compared to other bacteria. It is possible, for example, that no biofilm-dedicated transcriptional regulator(s) exist. In that scenario, if specific flux-dependent metabolites are allosteric regulators of enzymes that make or degrade adhesive mycomembrane components, aggregation and dispersal could be directly controlled by metabolic flux without a regulator serving as a middleman. Indeed, the absence of systems that affect biofilm formation in other bacteria, or the tenuous connection between these systems and biofilm formation in mycobacteria, may lend evidence in that direction. For example, quorum sensing systems affect biofilm formation in many bacteria, but there is no described quorum sensing system in mycobacteria. Similarly, there is an unclear role for c-di-GMP in NTM biofilms. Overexpressing an *E. coli* diguanylate cyclase in *M. smegmatis* increases mycolic acid synthesis and makes biofilms more robust, and knocking out the diguanylate cyclase/phosphodiesterase (MSMEG_2196) in *M. smegmatis* affects colony morphology(70, 71). However, bacteria that utilize c-di-GMP to regulate biofilms often encode dozens of enzymes that can produce (GGDEF domain) and/or degrade (EAL domain) the small molecule(72), and MSMEG_2196 is the only enzyme *M. smegmatis* encodes that processes c-di-GMP (it has both a GGDEF and EAL domain)(70). The utilization of guanosine for c-di-GMP production is also one of the ways in which purine biosynthesis is linked to biofilm formation in other bacterial species(73, 74). However, because supplemental adenosine and guanosine did not affect aggregation in either WT or Δ*purF,* our results indicate that in *M. smegmatis*, purine biosynthesis affects biofilm formation largely because of its utilization of glutamine.

The identification of glutamine as a flux-dependent sensor regulating NTM aggregation in response to nitrogen has precedence. While both glutamate and glutamine are amine donors for biosynthetic reactions, they play disparate roles in cellular physiology. The intracellular glutamate pool is very large as glutamate serves as a counterion to intracellular K^+^, and intracellular glutamate therefore fluctuates with external osmolarity(51). The large glutamate pool may also be important for driving transamination reactions, which have standard free energies near zero, forward(75). At least in enterics, intracellular glutamate concentrations do not change with nitrogen availability(51, 52). The intracellular glutamine pool is smaller and does swell and shrink in response to available nitrogen in a variety of bacteria(49, 50, 52). Our next steps are to determine exactly how glutamine impacts NTM matrix components. Glutamine directly interacts with nitrogen regulators like PII and GlnR, and NTM including *M. smegmatis* encodes both of those proteins. Indeed, an *M. abscessus* Δ*glnR* mutant has a biofilm defect, albeit with a concomitant defect in overall growth(76). It is also possible that glutamine affects another transcriptional regulator in NTM that directly affects matrix production, or that it directly affects an enzyme important for matrix production. We will also use the results of our transposon screen to further explore putative intracellular nodes that regulate NTM aggregation in response to carbon availability. For example, citrate, succinate, AKG, and PEP/pyruvate represent candidate nodes that we hit that are also well-documented flux-dependent sensors(29). Future studies will be carried out to identify the carbon node and determine how it interacts with glutamine, and how matrix production is ultimately affected.

NTM aggregation is intimately connected to metabolism, and we suspect that viewing NTM biofilm regulation through the lens of flux-dependent sensors could provide insight into the underlying regulatory circuits that have otherwise been difficult to achieve. By describing the core metabolic nodes that control aggregation, we hope to work outwards and determine how exactly those nodes mechanistically impact matrix/adhesion production. While *M. smegmatis* and *M. abscessus* seem to respond to chemical cues in similar ways in terms of aggregation regulation(26), it is not known if they share underlying biofilm regulatory systems. We will also use these *M. smegmatis* studies to launch parallel experimental strategies in *M. abscessus*, with the ultimate goal of revealing anti-biofilm therapeutic targets in pathogenic NTM.

## Methods and materials

### Strains, cloning, and growth conditions

Strains used in this study are listed in Table S2. Routine *M. smegmatis* cultures were grown at 37°C, shaking at 250 RPM in TYEM (10 g tryptone/L, 5 g <u>y</u>east <u>e</u>xtract/L, 2 mM <u>M</u>gSO_4_). Filter sterilized glucose, succinate, and glycerol, NH_4_Cl and NaNO_3,_ were added to sterile autoclaved TYEM when indicated. A modified M63 medium(26) (100 mM KH_2_PO_4_, 20 mM monosodium glutamate, 20 mM glucose, 0.5 mM proline, 1 mM MgSO_4_, 10 μM FeSO_4_, 1x SL-10 trace metal solution, and 20 mM NH_4_Cl [when noted], filter sterilized) was used as our defined medium. 15 mg/ml stock solutions of adenosine and guanosine solutions in DMSO were filter sterilized and added to 0.15 mg/mL where indicated. Recombineering was performed as described previously to create Δ*garA*, Δ*purF, and* Δ*gap* mutants in *M. smegmatis* mc^2^155, with minor modifications(26). 200-800 ng of linear DNA was used for the electroporation, and in the case of the Δ*purF* mutant, the chilled 10% glycerol wash used to make the cells electrocompetent was supplemented with 0.5 mM KoAC and 250 mM glucose. 500 bp flanking regions were amplified either by colony PCR or genomic extracts. After mutagenesis, the Δ*garA* mutant strain was cured of pJV53 by passaging on antibiotic free TYEM plates until it was verified as kanamycin sensitive. The Δ*gap* and Δ*purF* mutants were not cured of the pJV53 plasmid, so their WT comparator in the aggregation assays was WT + pJV53. Since pJV53 and our complement vector pSD5hsp60 both confer kanamycin resistance, a new complement vector, pSD5hsp60*hyg* was created to complement the Δ*purF* mutant. The pSD5hsp60 plasmid was linearized via PCR with primers that included homology with the sequence of the *hygR* cassette from pYUB1471. *hygR* and its promoter were amplified with primers that included homology to the pSD5hsp60 plasmid. The linearized pSD5pro vector and *hygR* fragment were then joined using NEBuilder HiFi DNA Assembly (NEB E2621L), transformed into chemically competent *E. coli* DH10B cells, and plasmids were isolated using Monarch plasmid miniprep kit (NEB T1010L). Plasmids were verified by full plasmid sequencing and transformed into electrocompetent *M. smegmatis* with the modifications to the electro-transformation method which are described above. Pellicle biofilms were diluted from TYEM cultures to .01 OD_600_ into 2 mL of M63 glucose medium in a well of a 24-well polystyrene plate and incubated statically at 37°C for 4 days prior to imaging.

### Aggregation assays

Aggregation assays were performed as described with minor modifications(26). Briefly, medium for aggregation assays was prepared in flasks and inoculated with the indicated strain of bacteria to an OD_600_ of 0.01. After inoculation, 5 mL aliquots were pipetted into disposable borosilicate culture tubes. These culture replicates were incubated at 37°C, shaking at 250 RPM. At each indicated timepoint, three culture replicates were harvested. For each replicate, the entire culture was poured through a 10 µm strainer. Culture that passed through the strainer was designated as the planktonic cell fraction, and the OD_600_ was immediately recorded. The original culture tube was washed with 5 mL of 1xPBS, which was then filtered through the aforementioned 10 µm strainer containing the aggregate fraction, to remove residual planktonic cells. Aggregates that remained on the strainer were resuspended in 4.5 mL 1xPBS + 6% Tween 20 and poured back in the original culture tube. 500 µL of Tween 20 was added for a final volume of 5 mL and a final Tween 20 concentration of 28.5%. Aggregate fractions were then water bath sonicated until no visible clumps remained, and the OD_600_ of the aggregate fraction was recorded.

### Transposon mutagenesis, colony morphology imaging, and arbitrarily primed PCR

#### Phage Titering and Propagation

50 µL of 10-fold dilutions of phiMycoMarT7 phage stock was added to 500 µL of WT *M. smegmatis* culture and allowed to infect for 15 minutes at room temperature. 4.5 mL of top agar (58.5 mL 7H9 media, 58.5 mL MBTA agar, 2 mM CaCl_2_) was added to each dilution, and the mixture was plated on 7H10 agar plates. After 2 days of growth at 30°C incubation, a titer was determined from plaque counts. To propagate the phage, 25 7H10 agar plates with ∼5000 plaques per plate were prepared using the titered phage stock to infect WT *M. smegmatis,* adding top agar as described in the titer step. When lacy lawns formed on the plates, 2.5 mL of phage buffer was added to each plate and placed on an orbital shaker at room temperature for 4 hours before being placed at 4°C overnight. The buffer and top agar were collected and spun down at 10,000 RPM for 10 minutes to pellet the agar. The supernatant was then filter sterilized. The phage titer process was then repeated to determine the concentration of the new stock.

#### Phage Transfection and primary colony morphology screen

WT *M. smegmatis* was grown in TYEM liquid culture, culture turbidity was quantified by OD_600_ reading, and the volume of culture needed for a final phage MOI of 10 was calculated. This volume of cells was pelleted by spinning for 7 minutes at 5000 RCF at 4°C. The supernatant was discarded, and the pellet was resuspended in 1 mL of 1x10^11^ PFU/mL phage stock. The cell-phage mixture was incubated at 37°C for 30 minutes before the mixture was added to 4 mL of 7H9+0.05% Tween 80 and incubated at 37°C with 250 RPM shaking for 3 hours. A 1 mL aliquot of the mixture was centrifuged at 5000 RCF for 1 minute and the supernatant was discarded. The pellet was resuspended in 1xPBS+0.05% Tween 80 and pelleted again by centrifuging at 5000 RCF for 1 minute. The supernatant was discarded, and the pellet was resuspended in 1 mL TYEM and plated onto a 15 mm x 150 mm TYEM agar plate supplemented with 40 µg/mL of kanamycin (Kan40). The plates were incubated at 37°C, and growth was observed at days 5 and 12. On each observation day, colonies displaying morphology different to WT were streaked onto smaller TYEM + Kan40 plates. This protocol was repeated for each for each of 14 ‘batches’ (designated with letters in mutant ID column of Table S1). Hits in the same gene in a single batch were considered redundant and were not listed as separate hits in Table S1.

#### Secondary Screen/Colony Morphology Imaging

Each selected mutant colony was cultured in TYEM, cultures were normalized to an OD_600_ of 1, and 4 µL of each sample was spotted onto TYEM agar plates alongside WT *M. smegmatis*. The plates were incubated at 37°C and imaged with a Zeiss Stereo Discovery V8 microscope at days 5 and 12. All TYEM colony morphology images shown in the paper are taken at day 5 unless noted otherwise. M63 colony morphology images were taken at day 2. Sterile paper discs were loaded with 10 µL of water, 5 µL of 15 mg/mL adenosine in DMSO and 15 mg/mL guanosine in DMSO, and 10 µL of DMSO, to a total volume of 10 µL for each condition.

#### Arbitrarily Primed PCR and Tn Mapping

500 µL liquid culture aliquots of each mutant candidate were heated at 95°C for 30 minutes, with vortexing every 5 minutes. PCR amplification of transposon insertion site was done in two rounds of PCR using a 50 µL reaction volume. PCR was performed using the Bio-Rad T100 thermocycler. PCR 1 consisted of 2 µL of boiled mutant candidate cell culture mixed with 25 µL OneTaq quick-load 2X master mix (NEB M0486L), 2.5 µL DMSO, 1 µL of 10 µM primer WD553, 1 µL of 10 µM primer WD557, and 18.5 µL ddH_2_O. The samples were then ran on PCR Cycle 1: Step 1 Initial Denaturing: 95°C for 5 minutes; 5 Cycles of Step 2: 95°C for 30 seconds, Step 3: 30°C for 30 seconds, Step 4: 72°C for 30 seconds; 30 Cycles of Step 6: 95°C for 30 seconds, Step 7: 38°C for 30 seconds, Step 8: 72°C for 30 seconds; Final Extension: 72°C for 7 minutes. For each PCR 1 reaction, 10 µL of PCR product sample was combined with 25 µL OneTaq quick-load 2X master mix, 2.5 µL DMSO, 1 µL of 10 µM primer WD414, 1 µL of 10 µM primer WD555, and 10.5 µL ddH_2_O. PCR was performed as above, but with the Cycle 2 settings: Step 1 Initial Denaturing: 94°C for 30 seconds; 30 Cycles of Step 2: 94°C for 30 seconds, Step 3: 56°C for 30 seconds, Step 4: 68°C for 30 seconds; Final Extension: 68°C for 7 minutes. PCR product size was verified by gel-electrophoresis. The PCR 2 products were purified using the Monarch PCR & DNA Cleanup Kit (NEB #T1030) according to the manufacturer’s instructions, with a final elution volume of 10 µL of ddH_2_O. The concentration of each mutant sample was quantified using a NanoDrop One spectrophotometer. 10 ng/µL of PCR product was combined with 25 pmol/µL of primer WD554, brought to a total volume of 15 µL, and sent to Azenta for Sanger sequencing. The sequencing results were compared against the reference genome for WT *M. smegmatis* MC^2^155 using a BLAST search and the Tn insertion site was determined.

### Targeted metabolomics

Metabolite extraction was performed as described with modifications(32). *M. smegmatis* culture was grown in M63 defined medium as described in the Results section. For each experimental condition, five replicate cultures were set up per time point. Each replicate contained a total of 15 mL of cell culture. At each indicated time point, all 15 mL of culture was poured through a Durapore 0.22 µm PVDF Membrane Filter (MilliporeSigma). The membrane was then washed with 5 mL of 1xPBS, pre-heated to 37°C. The filter membrane was immediately moved to 2 mL of dry ice-chilled 2:2:1 acetonitrile:methanol:water solution for cell lysis. The cells were scraped off the filter membrane with a cell scraper and collected into a 2 mL bead beating tube from the Precellys Tough Micro-organism Lysing Kit VK05 (Bertin Instruments). The cells were bead-beaten with an Omni International Bead Ruptor 12 machine (4 speed, 3 cycles, 2 minutes per cycle, 10 seconds break in between cycles; all done at 4°C). The resulting cell lysate was pelleted in a microcentrifuge at 20,000 RCF for 5 min at 4°C. From the supernatant, a 600 µL aliquot was transferred to a Costar Spin-X 0.22 µm Centrifuge Tube Filter (Corning), and the rest was decanted. The supernatant was passed through the filter in a microcentrifuge using the same setting, and the resulting 600 µL metabolite extract was immediately capped and frozen at -80°C until metabolomic analysis by LC-HRMS..

The remaining cell lysate pellet was set on a heat block in a chemical fume hood at 55°C for 2 hours to evaporate any remaining acetonitrile:methanol:water solution. Once dried, the pellet was resuspended in 500 μL of 3% SDS in 10 mM Tris-HCl solution, pH 7.5. The resuspended pellet was bead-beaten again in the same manner as described, and the resulting liquid was left in a 4°C cold room to settle for 20 minutes. From the settled liquid, a 1:10 dilution in water was made and the protein concentration within the cell lysate was assessed using the Pierce™ Dilution-Free™ Rapid Gold BCA Protein Assay (Thermo ScientificTM).

Targeted metabolomics analysis was done by the University of Pittsburgh Health Sciences Mass Spectrometry Core(MSC). Extracted metabolite samples were sent to the MSC, where the intracellular pool of glutamine was quantified using LC-HRMS. Briefly, a deuterated internal standard mix that included creatinine-d_3_, alanine-d_3_, taurine-d_4_ and lactate-d_3_ (Sigma-Aldrich) was added to the sample lysates at a final concentration of 10 µM. After 3 minutes of vortexing, the supernatant was cleared of protein by centrifugation at 16,000xg. Cleared supernatant (2 µL) was injected via a Thermo Vanquish UHPLC and separated over a reversed phase Thermo HyperCarb porous graphite column (2.1×100 mm, 3 μm particle size) maintained at 55°C. For the 20-minute LC gradient, the mobile phase consisted of the following: solvent A (water/0.1% FA) and solvent B (ACN/0.1% FA). The gradient was the following: 1%B for the first minute increasing to 15%B over 5 minutes, followed by an increase to 98%B over 5 minutes that was held for 5 minutes before equilibration at initial conditions for 5 minutes. The Thermo ID-X tribrid mass spectrometer was operated in both positive and negative ion mode, scanning in ddMS^2^ mode (2 μscans) from 70 to 800 *m/z* at 120,000 resolution with an AGC target of 2e5 for full scan, and 2e4 for MS^2^ scans using HCD fragmentation at stepped 15,35,50 collision energies. Source ionization was 3.0 and 2.4kV spray voltage respectively, for positive and negative mode. Source gas parameters were 35 sheath gas, 12 auxiliary gas at 320°C, and 8 sweep gas. Calibration was performed prior to analysis using the Pierce^TM^ FlexMix Ion Calibration Solution (Thermo Fisher Scientific). Integrated peak areas were extracted manually using Quan Browser (Thermo Fisher Xcalibur ver. 2.7). Targeted metabolite values are reported as the ratio of the analyte to the internal standard before conversion to absolute concentration via calibration curves ranging from 15 fmol/µL to 100 pmol/µL.

### Statistical Analysis

Differences between metabolite averages were analyzed using Student’s t-test for comparisons between two groups. One-way ANOVA and post hoc Tukey-Kramer multiple comparison test was used for comparisons between three or more groups. (*) indicates statistical significance with a *p*-value < 0.05; (**) indicates statistical significance with a *p*-value < 0.01. All analysis was performed in GraphPad Prism 10.

## Acknowledgements

This work was supported by the NIH (NIAID R01-AI170607 to W.H.D. and S10-OD032141 to S.L.G.) and the Cystic Fibrosis Foundation (BOMBER21R3 to W.H.D.). M.M. and Y.W. were supported by an RK Mellon Institute for Pediatric Research Trainee Award and a Children’s Hospital of Pittsburgh of UPMC Research Advisory Committee Graduate Student Fellowship, respectively. Work performed in the Health Sciences Mass Spectrometry Core (RRID:SCR_025222) and services and instruments used in this project were graciously supported, in part, by the University of Pittsburgh and the Office of the Senior Vice Chancellor for Health Sciences. Research was conducted at the University of Pittsburgh John G. Rangos Sr. Research Center and the University of Pittsburgh Health Sciences Mass Spectrometry Core. Thanks to Graham Hatfull, Katherine Wetzel, and Eric Rubin for providing phiMycoMarT7 and for advice and protocols germane to the transposon mutagenesis screen.

